# Correlation between increased atrial expression of genes related to fatty acid metabolism and autophagy in patients with chronic atrial fibrillation

**DOI:** 10.1101/815126

**Authors:** Yasushige Shingu, Shingo Takada, Takashi Yokota, Ryosuke Shirakawa, Akira Yamada, Tomonori Ooka, Hiroki Katoh, Suguru Kubota, Yoshiro Matsui

## Abstract

Atrial metabolic disturbance contributes to the onset and development of atrial fibrillation (AF). Autophagy plays a role in maintaining the cellular energy balance. We examined whether the altered atrial expression of genes related to fatty acid metabolism is linked to that related to autophagy in chronic AF. Right atrial tissue was obtained during heart surgery from 51 patients with sinus rhythm (SR, n=38) or chronic AF (n=13). Preoperative fasting serum free-fatty-acid levels were significantly higher in the AF patients. The atrial gene expression of fatty acid binding protein 3 (*FABP3*), which is involved in the cells’ fatty acid uptake and intracellular fatty acid transport, was significantly increased in AF patients compared to SR patients; in the SR patients it was positively correlated with the right atrial diameter and intra-atrial EMD, parameters of structural and electrical atrial remodeling that was evaluated by an echocardiography. In contrast, the two groups’ atrial contents of diacylglycerol (DAG), a toxic fatty acid metabolite, were comparable. Importantly, the atrial gene expression of microtubule-associated protein light chain 3 (*LC3*) was significantly increased in the AF patients, and autophagy-related genes including *LC3* were positively correlated with the atrial expression of *FABP3*. In conclusion, in chronic AF patients, the atrial expression of *FABP3* was upregulated in association with autophagy-related genes without altered atrial DAG content. Our findings may support the hypothesis that dysregulated cardiac fatty acid metabolism contributes to the progression of AF and induction of autophagy has a cardioprotective effect against cardiac lipotoxicity in chronic AF.

## Introduction

Atrial fibrillation (AF) is the most common cardiac arrhythmia, and its presence is associated with increased risks of death, heart failure, and stroke [1–3]. With the current increased prevalence of AF, the prevention of AF is important not only for public health but also to reduce the associated economic burden [4]. The risk factors for AF are diverse, including higher serum levels of free fatty acids (FFAs), obesity, hypertension, inflammation, and oxidative stress [5–7]. The mechanisms underlying the onset and development of AF have not been fully elucidated.

The pathophysiology of AF is complex and involves electrical, structural, contractile, and neurohormonal remodeling [8, 9]; metabolic disturbance in the atrial cardiac muscle is a recent focus of AF research, as the heart has a very high energy demand due to its organ-specific feature involving the constant activation of mitochondrial oxidative phosphorylation. In particular, fatty acids are the major fuel for the heart; their use depends on their uptake into the cells, transport from the cytosol into the mitochondria, and β-oxidation in the mitochondria. Prior research has shown that an elevated level of circulating FFAs is a strong risk factor for AF and AF-related stroke [6, 10] and can be a trigger of cardiac lipotoxicity, which is defined as the excess accumulation of toxic fatty acid metabolites such as diacylglycerol (DAG) in the heart. This may occur when the influx of FFAs exceeds the intracellular fatty acid oxidation, which leads to cardiac dysfunction, cardiac remodeling, and arrhythmias [11]. However, it is still unclear how metabolic disturbances including abnormal fatty acid metabolism contribute to the development of AF.

Autophagy, the process of the degradation of intracellular components (e.g., proteins) in lysosomes, plays an important role in cellular homeostasis via cellular quality control. Autophagy was also shown to contribute to the cellular energy balance, in particular through a mechanism of fatty acid metabolism termed “lipophagy” (the degradation of excess lipids by autophagy) and the degradation of lipid stores in the cells [12]. Accordingly, autophagy may regulate the fatty acid metabolism in cardiomyocytes. Alterations of the autophagy in the atrial muscles of patients with persistent AF [13, 14] or post-operative AF have been reported [15]. Although it is still controversial whether the induction of autophagy has a cardioprotective or detrimental effect in AF, it is possible that autophagy is involved in metabolic remodeling in the atrium in chronic AF patients.

We conducted the present study to determine: (1) whether the expression of genes related to fatty acid metabolism and autophagy are altered in the atria of patients with chronic AF, and (2) whether changes in these gene expression patterns are correlated with each other. We used human atrial tissue excised from patients during cardiac surgery, and our findings provide new insight into the pathophysiology of AF, focusing on fatty acid metabolism and autophagy in the human atrium.

## Materials and Methods

### Patients

This study was conducted at Hokkaido University Hospital and Teine Keijinkai Hospital and included 51 consecutive patients: 38 with sinus rhythm (SR) and 13 with chronic AF who underwent cardiovascular surgery between 2013 and 2019 at either of these hospitals. All of the patients were Japanese. The patients with SR in the present series partly overlap with those of our recently published report [16].

After the establishment of a cardiopulmonary bypass (10 min after the infusion of heparin 300 IU/kg), right atrial myocardial tissue (approx. 100 mm^2^) was excised from the right atrial incision site or the insertion point of a drainage cannula. The tissue was frozen and stored at −80°C until analysis.

Type 2 diabetes was defined as a fasting glucose level ≥7.0 mmol/L and/or taking antidiabetes medications. Coronary artery disease was evaluated by coronary angiography, and stenosis ≥75% was defined as significant; a patient with a history of percutaneous coronary intervention was also regarded as having coronary artery disease.

The study protocol was approved by the Ethics Committees of Hokkaido University Hospital and Teine Keijinkai Hospital and performed according to the Declaration of Helsinki. Written informed consent was obtained from each patient before the surgery. This study was registered in the UMIN Clinical Trials Registry: UMIN000012405 and UMIN000018137.

### Transthoracic echocardiography

A Vivid Seven system (GE/Vingmed, Milwaukee, WI, USA) with an M3S (2.5–3.5 MHz) transducer, an Aplio system (Toshiba Medical Systems, Tokyo) with a 2.5-MHz transducer, or a Philips system (Philips Ultrasound, Bothell, WA) with a 2.5-MHz transducer was used for the pre-operative echocardiography. The left ventricular and atrial diameters were measured from the parasternal long-axis view. The right atrial diameter (the largest minor-axis diameter in the four-chamber view) was obtained at end-systole [17].

The electromechanical delay (EMD) of the right atrium was assessed in all but one of the SR patients as a time interval (T1) from the beginning of the P-wave on surface ECG to the beginning of the late diastolic wave (A’) of the tricuspid annulus [17]. T2 was the time interval from the beginning of the P-wave on the surface ECG to the A’ of the septal mitral annulus. The intra-atrial EMD was the interval from T1 to T2, expressed as T2-T1 (msec). The left ventricular ejection fraction (LVEF) was measured using the biplane method of disks. The mitral regurgitation grade was defined by the regurgitation jet area-to-left atrium ratio (mild, <20%; moderate, 20%–40%; severe, >40%).

Tricuspid regurgitation was graded using the regurgitation jet area (mild, <5 cm^2^; moderate, 5–10 cm^2^; severe, >10 cm^2^). Aortic stenosis was defined using the valve area (mild, >1.5 cm^2^; moderate, 1.0–1.5 cm^2^; severe, <1.0 cm^2^). Aortic regurgitation was determined using a combination of the jet width/outflow tract, the pressure half-time, and the diastolic reverse flow at the abdominal aorta (mild, moderate, severe) [18].

### Blood biochemistry

Blood was collected from each patient early in the morning after a 10-hr fast within 2 days before surgery. The blood glucose, hemoglobin A1c, insulin, and FFA levels were measured by an enzymatic reaction involving glucose oxidase, high-performance liquid chromatography, an enzyme immunoassay, and the enzymatic method, respectively. Total cholesterol and triglycerides were assessed by enzymatic methods. The plasma levels of B-type natriuretic peptide (BNP) were evaluated by a chemiluminescent enzyme immunoassay.

### The atrial expression of genes related to fatty acid metabolism and autophagy

The atrial expression of the following genes related to fatty acid metabolism and autophagy was determined by reverse transcription-polymerase chain reaction (RT-PCR): cluster of differentiation 36/fatty acid translocase (*CD36*), carnitine palmitoyltransferase 1B (*CPT1B*), fatty acid-binding protein 3 (*FABP3*), autophagy-related gene 5 (*ATG5*), Unc-51-like kinase 1 (*ULK1*), Beclin-1 (*BCLN1*), and microtubule-associated protein light chain 3 (*LC3*). Myocardial mRNA was isolated from frozen tissue using a High Pure RNA Tissue Kit (Roche, Penzberg, Germany) and was then reverse-transcribed into cDNA using a Transcriptor First Strand cDNA Synthesis Kit (Roche).

The RT-PCR was performed using FastStart Essential DNA Probes Master (Roche) and the Real-time Ready Assay (Roche Assay ID: *CD36,* 144833; *CPT1B*, 126501; *FABP3*, 118811; *ATG5*, 125999; *ULK1*, 109914; *BCLN1*, 100115; *LC3*, 144582; *TBP*, 143707). PCR amplification was then performed with a reaction volume of 20 μL using a LightCycler Nano (Roche) under the conditions specified by the manufacturer. After the initial denaturation and activation of the enzyme for 10 min at 95°C, 45 cycles of denaturation at 95°C for 10 sec, and annealing and extension at 60°C for 30 sec were performed. *TBP* (TATA-binding protein) was used as a reference gene, as we confirmed that TBP is not influenced by the occurrence of AF.

### The atrial enzymatic activities of the mitochondrial TCA cycle and fatty acid β-oxidation

We spectrophotometrically determined the activity levels of both citrate synthase (CS), a key enzyme in the tricarboxylic acid (TCA) cycle, and β-hydroxyacyl CoA dehydrogenase (β-HAD), a key enzyme in fatty acid β-oxidation, in the myocardial samples as described [19]. Due to the lack of atrial muscle samples, these enzymatic activities were measured in 20 of the 38 patients with SR and eight of the 13 patients with AF.

### The atrial DAG content

Heart tissue was homogenized in 1.5 mL of methanol, followed by mixing with 2.25 mL of 1 M NaCl and 2.5 mL of chloroform. The mixture was centrifuged at 1,500 g for 10 min at 4°C, and the organic phase was dried. DAG was then detected with a DAG assay kit (Cell Biolabs, San Diego, CA) following the manufacturer’s instructions. Due to the lack of atrial muscle samples, the atrial DAG content was measured in six patients with SR and six patients with AF.

### Statistical analyses

Values are presented as the mean ± standard deviation (SD) or the median (interquartile range [IQR]) as appropriate. We used Student’s t-test for continuous variables that are normally distributed and the Mann-Whitney U-test for other continuous variables. The chi-square test or Fisher’s exact test were used for categorical variables. We conducted a Pearson’s correlation analysis to determine linear relationships between continuous variables. The statistical analyses were performed using GraphPad Prism ver. 8 (GraphPad Software, San Diego, CA), and significance was defined as p<0.05.

## Results

### Patient characteristics

The characteristics of the SR and AF patients are summarized in **Table 1**. The median duration of AF was 12 years (range 1–16 years). Except for the use of medications, all parameters including cardiovascular risk factors were comparable between the two groups. The surgical procedures performed after the excision of atrial specimens included aortic valve replacement in 30 cases, total arch replacement in eight cases, coronary artery bypass grafting in 10 cases, aortic root replacement in six cases, and mitral valve repair in 12 cases. Multiple procedures were performed in some patients.

**Table 1.** Characteristics of the SR and AF patients.

|  | **SR**<br>**(n=38)** | **AF**<br>**(n=13)** | **p-value** |
| --- | --- | --- | --- |
| Age, yrs | 70 ± 13 | 69 ± 9 | 0.775 |
| Male | 18 (47%) | 8 (62%) | 0.378 |
| BMI, kg/m<sup>2</sup> | 22.8 ± 3.4 | 24.8 ± 4.3 | 0.097 |
| Heart rate, bpm | 67 ± 10 | 73 ± 13 | 0.134 |
| Systolic blood pressure, mm Hg | 118 ± 19 | 115 ± 18 | 0.643 |
| Diastolic blood pressure, mm Hg | 60 ± 12 | 68 ± 13 | 0.051 |
| Diabetes mellitus | 7 (18%) | 2 (15%) | 0.804 |
| Coronary artery disease | 13 (34%) | 2 (15%) | 0.198 |
| Medications |  |  |  |
| Diuretics | 18 (47%) | 13 (100%) | 0.001 |
| β-blockers | 14 (37%) | 9 (69%) | 0.043 |
| Statins | 18 (47%) | 4 (31%) | 0.297 |
| Total cholesterol, mmol/L | 4.7 ± 0.9 | 4.4 ± 0.7 | 0.357 |
| Triglycerides, mmol/L | 1.4 ± 0.7 | 1.0 ± 0.5 | 0.052 |
| Fasting blood glucose, mmol/L | 5.5 (0.8) | 5.7 (0.7) | 0.802 |
| Insulin, μU/mL | 5.0 (4.0) | 5.8 (8.9) | 0.612 |
| HbA1c, % | 5.7 ± 0.8 | 5.9 ± 0.6 | 0.324 |
| BNP, pg/mL | 79 (270) | 249 (321) | 0.060 |

Data are mean±SD, median (interquartile range), or n (%). AF, atrial fibrillation; BMI, body mass index; BNP, B-type natriuretic peptide; HbA1c, hemoglobin A1c; SR, sinus rhythm.

### Transthoracic echocardiography

As shown in **Table 2**, the diameters of the left and the right atria were significantly increased in the AF group. The left ventricular size (i.e., left ventricular end-diastolic diameter [LVDd]) and the left ventricular systolic function (i.e., the LVEF) were comparable between the two groups. Regarding valvular heart disease, the grade of mitral regurgitation was higher in the AF group than in the SR group.

**Table 2.** Echocardiographic parameters of the SR and AF patients.

|  | SR<br>(n=38) | AF<br>(n=13) | p-value |
| --- | --- | --- | --- |
| LVDd, mm | 48 (20) | 59 (17) | 0.209 |
| LVDs, mm | 31 (22) | 41 (20) | 0.132 |
| LVEF, % | 59 ± 14 | 54 ± 14 | 0.268 |
| Left atrial diameter, mm | 41 ± 7 | 53 ± 9 | <0.001 |
| Right atrial diameter, mm | 34 ± 6 | 42 ± 9 | 0.007 |
| Aortic stenosis: |  |  | 0.130 |
| Mild | 1 (3%) | 0 (0%) |  |
| Moderate | 1 (3%) | 1 (8%) |  |
| Severe | 15 (39%) | 1 (8%) |  |
| Aortic regurgitation: |  |  | 0.306 |
| Mild | 18 (47%) | 4 (31%) |  |
| Moderate | 4 (11%) | 2 (15%) |  |
| Severe | 10 (26%) | 2 (15%) |  |
| Mitral regurgitation: |  |  | 0.002 |
| Mild | 29 (76%) | 5 (38%) |  |
| Moderate | 2 (5%) | 5 (38%) |  |
| Severe | 2 (5%) | 3 (23%) |  |
| Tricuspid regurgitation: |  |  | 0.133 |
| Mild | 26 (68%) | 6 (46%) |  |
| Moderate | 3 (78%) | 3 (23%) |  |
| Severe | 0 (0%) | 1 (8%) |  |

Data are mean±SD, median (interquartile range), or n (%). AF, atrial fibrillation; LVDd, left ventricular end-diastolic diameter; LVDs, left ventricular end-systolic diameter; LVEF, left ventricular ejection fraction; SR, sinus rhythm.

### The serum levels of FFA and the atrial expression of genes related to fatty acid metabolism

The serum FFA levels were significantly higher in the AF group compared to the SR group (719 ± 107 vs. 416 ± 37 μmol/L, p=0.001, **Fig 1**).

**Fig 1:**
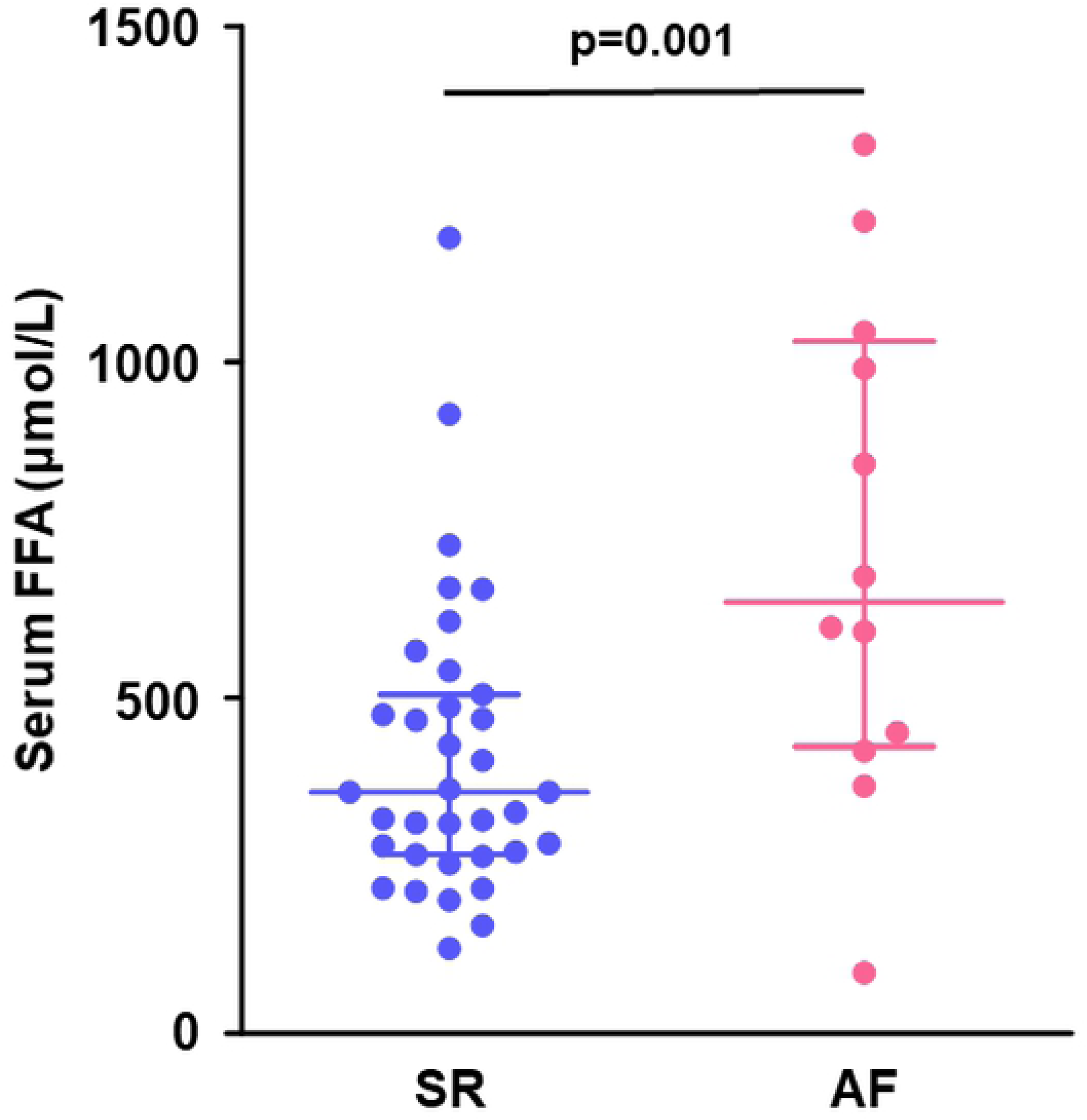
Preoperative levels of serum FFA. Lines indicate the median with the interquartile range (IQR) in each group (SR, n=35; AF, n=12). AF, atrial fibrillation; FFA, free fatty acids; SR, sinus rhythm.

The expressions of genes related to fatty acid metabolism in the right atrial muscle are shown in **Fig 2**. The gene expression of *CD36*, which facilitates fatty acid uptake across the plasma membrane, tended to be higher in the AF group than in the SR group (**Fig 2A**). The gene expression of *FABP3*, which facilitates fatty acid uptake into the cell and intracellular fatty acid transport, was significantly increased in the AF group (**Fig 2B**), but there was no significant difference between the two groups in the atrial expression of *CPT1B*, which is located on the outer mitochondrial membrane for fatty acid transport into the mitochondria (**Fig 2C**).

**Fig 2:**
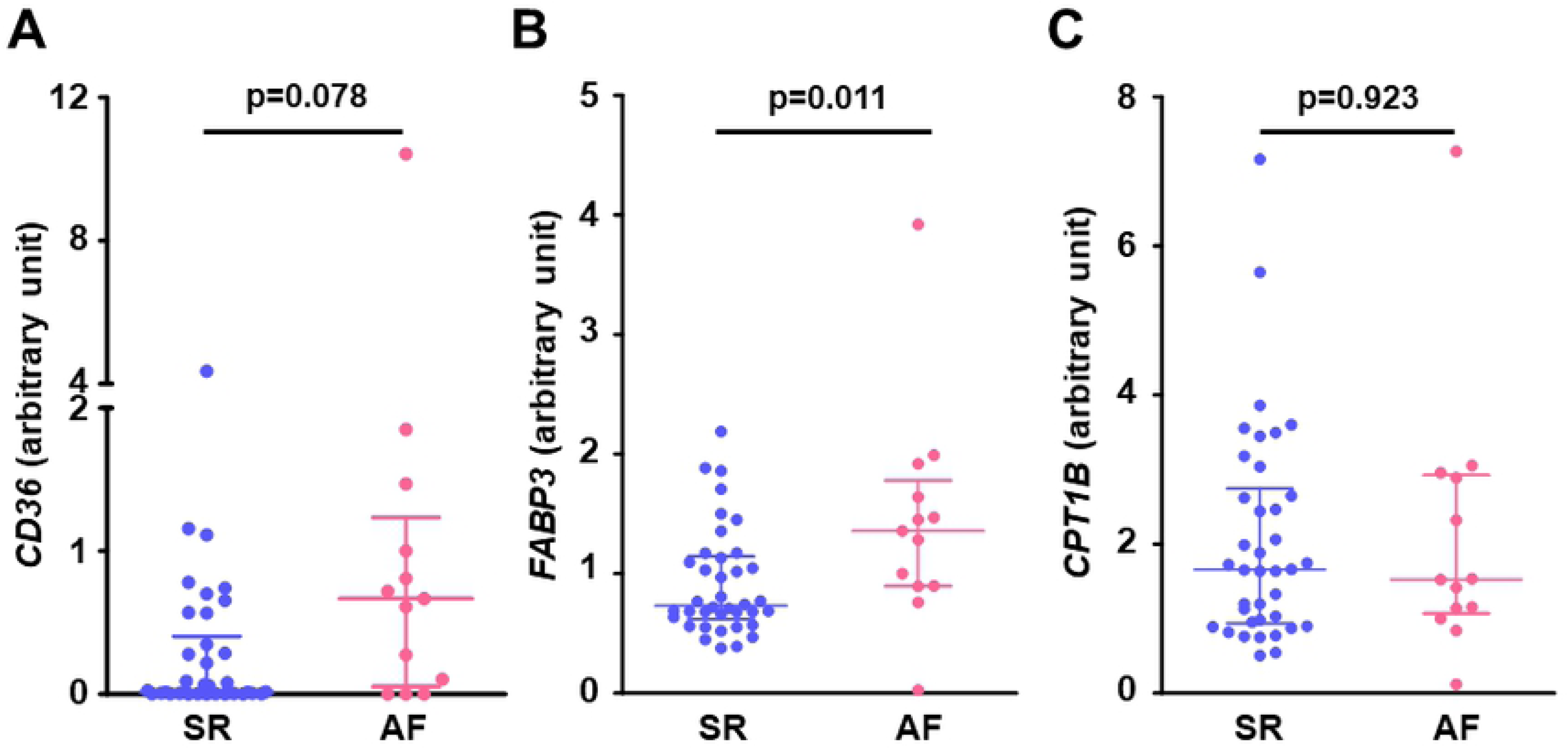
Gene expression related to fatty acid metabolism in the right atrial myocardium. (A) *CD36*, (B) *FABP3*, and (C) *CPT1B*. Lines indicate the median with IQR in each group (SR; n=38, AF; n=13). *CD36*, cluster of differentiation 36 (fatty acid translocase); *CPT1B*, carnitine palmitoyltransferase 1B; *FABP3*, fatty acid binding protein 3.

In the SR patients, *FABP3* gene was positively correlated with the right atrial diameter (**Fig 3A**) and the intra-atrial EMD (**Fig 3B**), indicating that increased atrial gene expression related to intracellular fatty acid transport was associated with structural and electrical atrial remodeling.

**Fig 3:**
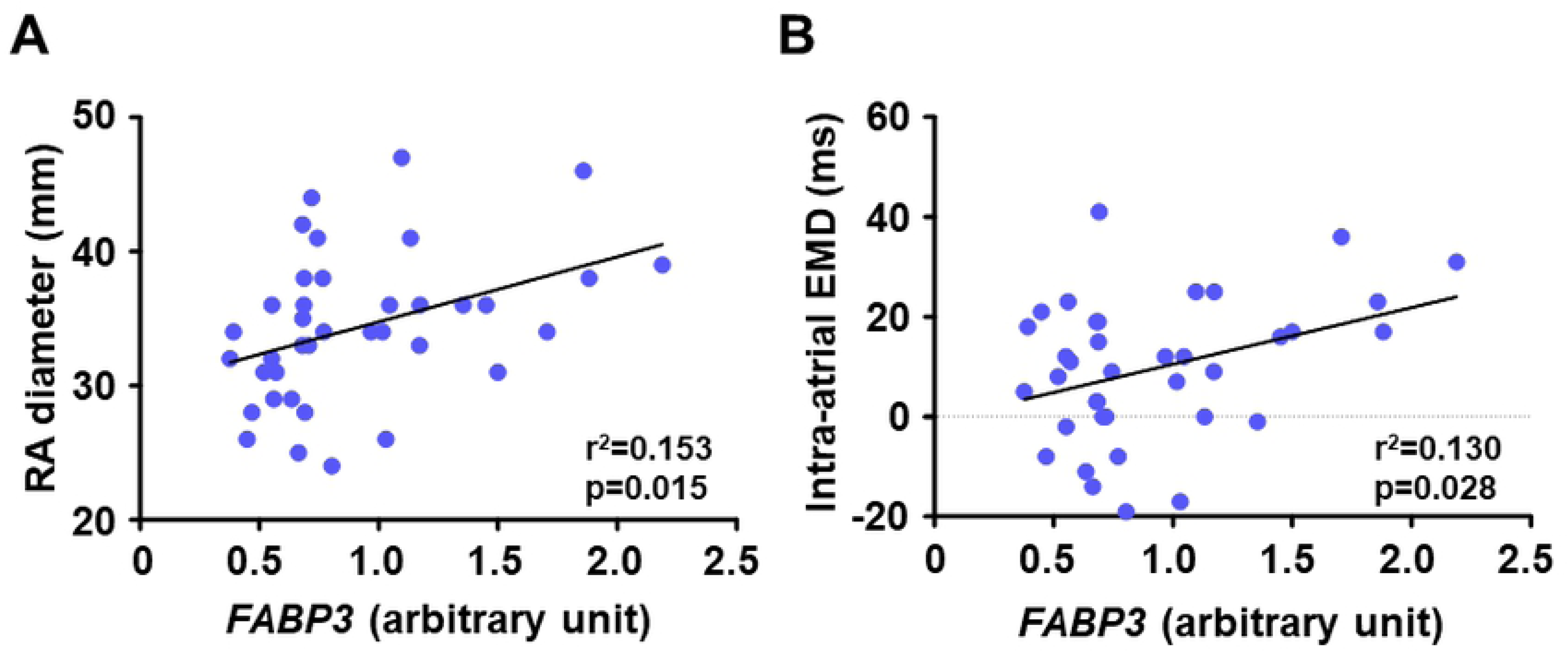
Association between gene expression levels of *FABP3* in the right atrial myocardium and parameters of atrial remodelling in patients with SR. (A) *FABP3* and RA diameter (n=38) and (B) *FABP3* and intra-atrial EMD (n=37). EMD, electromechanical delay; *FABP3*, fatty acid binding protein 3; RA, right atrium.

### The atrial enzymatic activities of the mitochondrial TCA cycle and fatty acid β-oxidation

Fig 4 illustrates the enzymatic activities related to the mitochondrial fatty acid β-oxidation and TCA cycle in the right atrial muscle. The CS activity was comparable between the SR and AF groups (Fig 4A), as was the β-HAD activity (Fig 4B).

**Fig 4:**
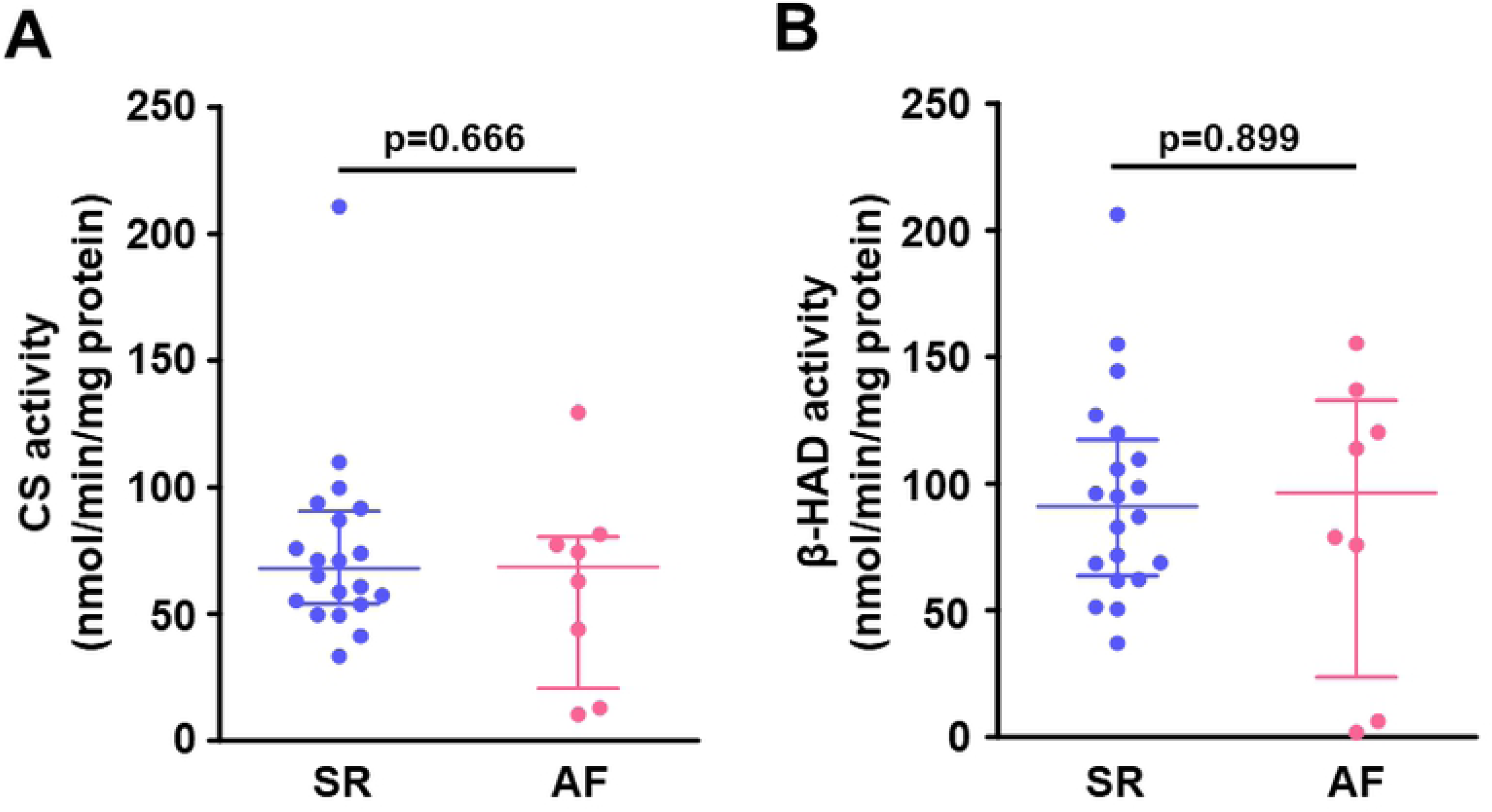
Enzymatic activities related to the mitochondrial TCA cycle and fatty acid β-oxidation in the right atrial myocardium. (A) CS activity and (B) β-HAD activity. Lines indicate the median with IQR in each group (SR, n=20; AF, n=8). β-HAD, β-hydroxyacyl CoA dehydrogenase; CS, citrate synthase.

### The atrial DAG content

Despite the higher serum levels of FFA and the upregulated atrial gene expression of *FABP3* in the AF group, there was no significant difference in the atrial content of DAG, a major toxic fatty acid metabolite, between the SR and AF groups (Fig 5).

**Fig 5:**
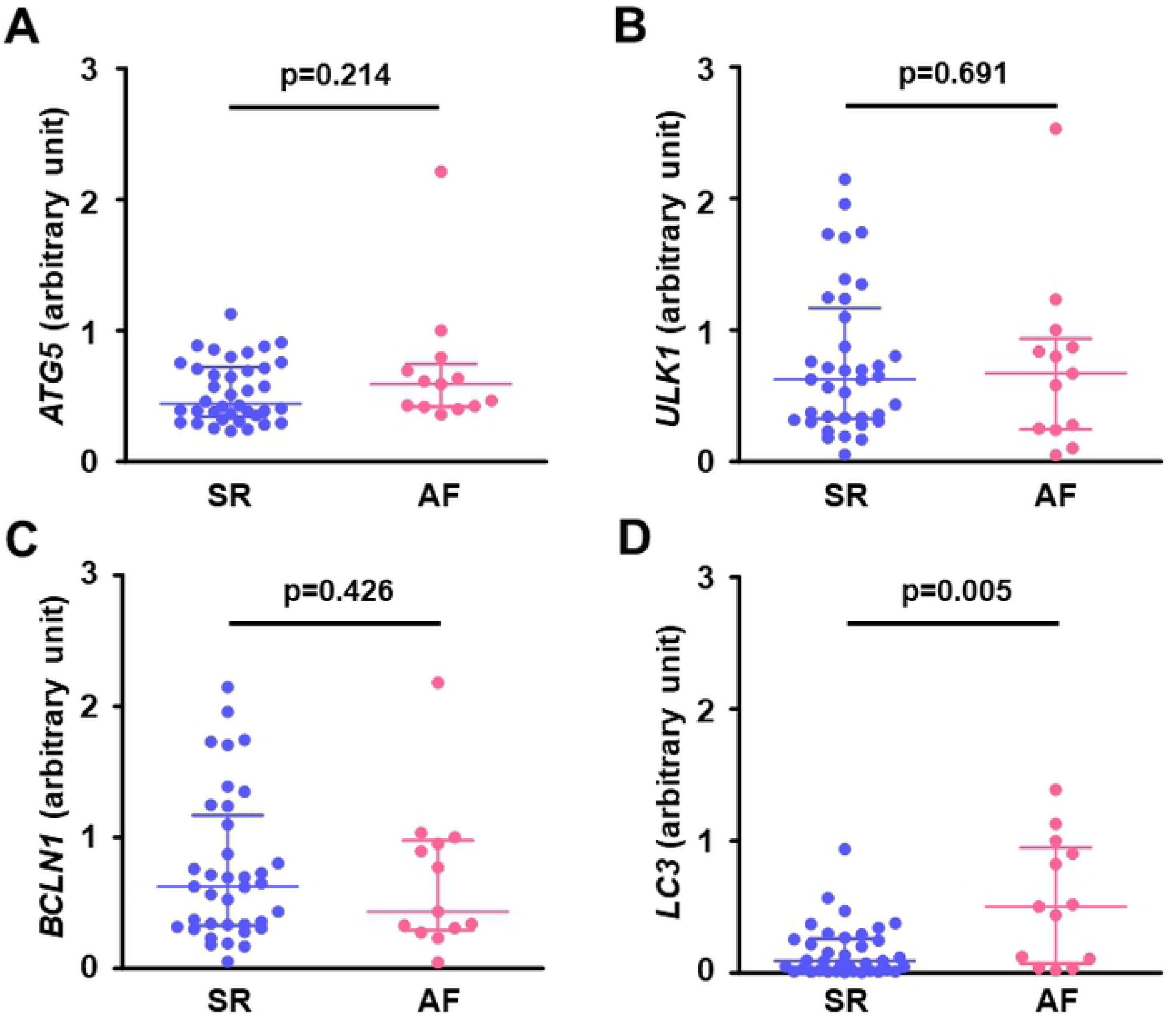
DAG contents in the right atrial myocardium. Lines indicate the median with IQR in each group (SR, n=6; AF, n=6). DAG, diacylglycerol; RFUs, relative fluorescence units.

### The atrial expression of autophagy-related genes

The gene expression of *LC3* in the right atrial muscle was significantly higher in the AF group than in the SR group (Fig 6D), but there was no significant difference in other autophagy-related genes including *ATG5*, *ULK1*, and *BCLN1* between the SR and AF groups (Figs 6A–6C).

**Fig 6:**
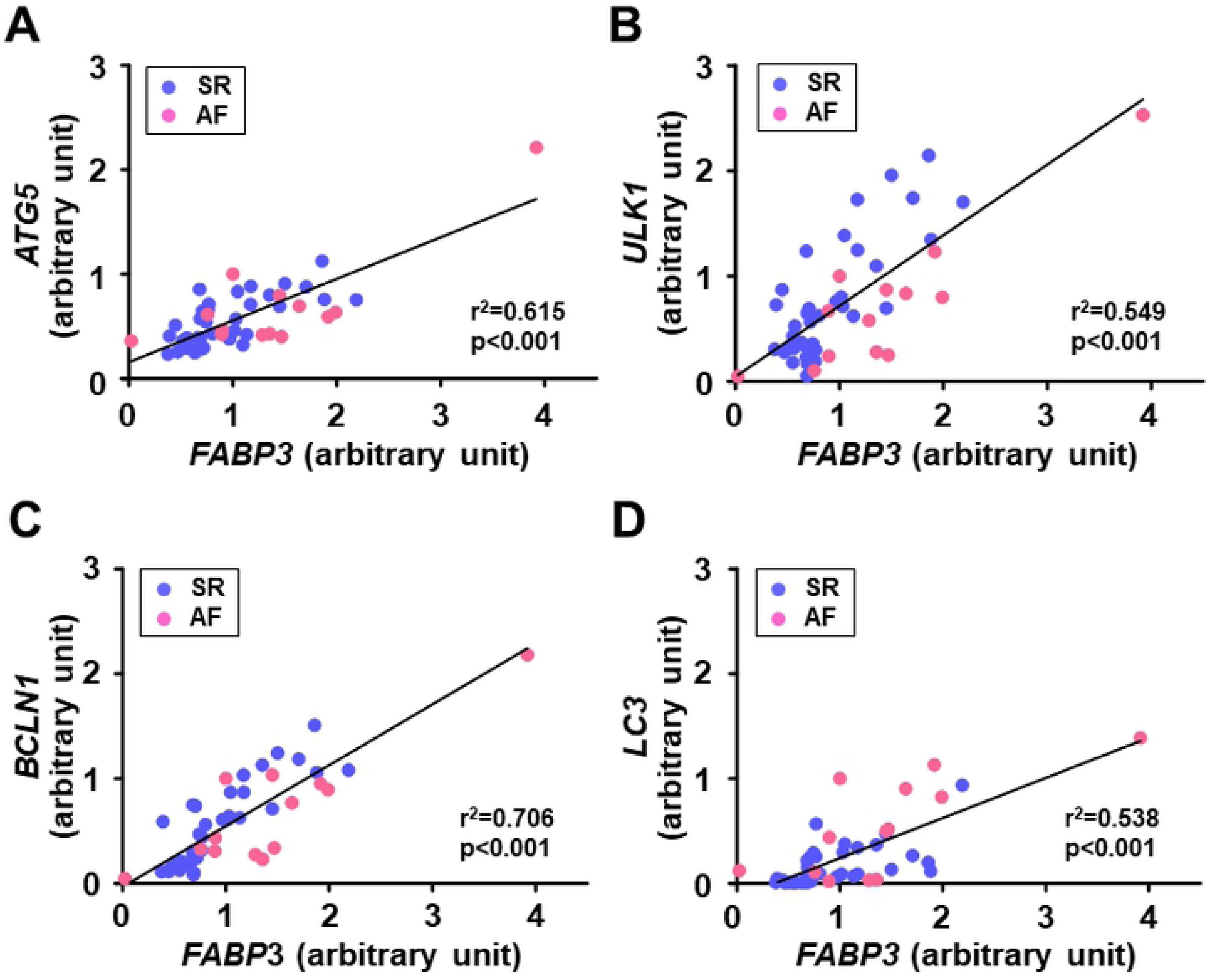
Gene expression related to autophagy in the right atrial myocardium. (A) *ATG5*, (B) *ULK1*, (C) *BCLN1*, and (D) *LC3*. Lines indicate the median with IQR in each group (SR, n=37 except for *ATG5* [n=38]; AF, n=13). *ATG5*, autophagy-related gene 5; *BCLN1*, beclin-1; *LC3*, microtubule-associated protein light chain 3; *ULK1*, Unc-51-like kinase 1.

### The linear relationship between the atrial expression of FABP3 gene and autophagy-related genes

The gene expression of *FABP3* was positively correlated with autophagy-related genes including *LC3* gene in all patients (Fig 7). Similar correlations were observed between *FABP3* gene and autophagy-related genes even when they were analyzed separately for the SR group (*FABP3* and *ATG5*: r^2^=0.510, p<0.001; *FABP3* and *ULK1*: r^2^=0.622, p<0.001; *FABP3* and *BCLN1*: r^2^=0.740, p<0.001; *FABP3* and *LC3*: r^2^=0.384, p<0.001) and the AF group (*FABP3* and *ATG5*: r^2^=0.670, p<0.001; *FABP3* and *ULK1*: r^2^=0.790, p<0.001; *FABP3* and *BCLN1*: r^2^=0.768, p<0.001; *FABP3* and *LC3*: r^2^=0.534, p=0.005).

**Fig 7:**
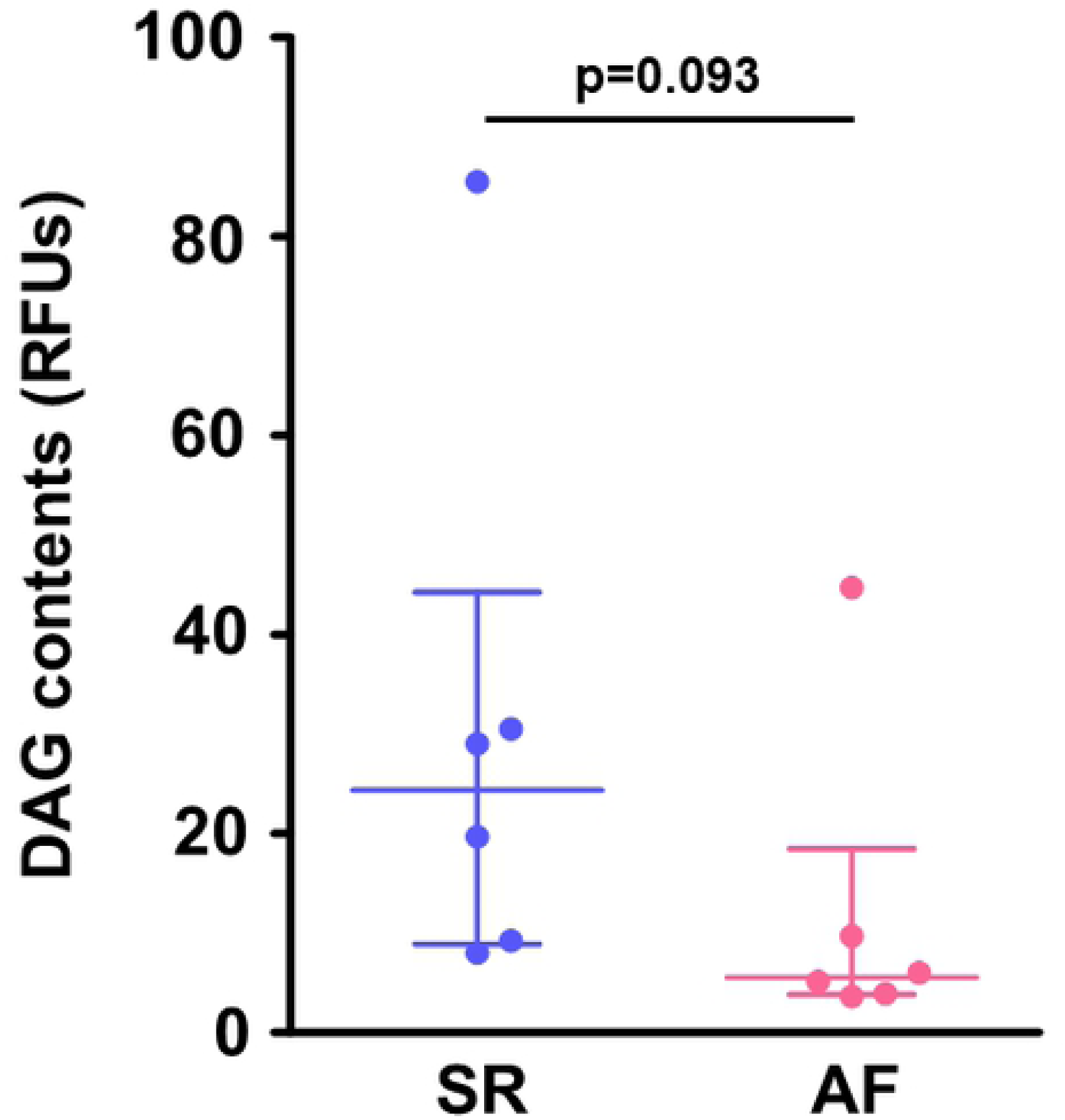
Association between the expression levels of *FABP3* gene and autophagy-related genes in the right atrial myocardium in patients with SR or AF. (A) *FABP3* and *ATG5* (SR; n=13; AF, n =38), (B) *FABP3* and *ULK1* (SR, n=13; AF, n=37), (C) *FABP3* and *BCLN1* (SR, n=13; AF, n=37), and (D) *FABP3* and *LC3* (SR, n=13; AF, n=37). *FABP3*, fatty acid binding protein 3. Other abbreviations are explained in the Fig 6 legend.

## Discussion

Our findings demonstrated that the atrial expression of a gene involved in fatty acid metabolism, *FABP3*, was upregulated in patients with chronic AF compared to the SR patients. In the SR patients, the increased atrial expression of *FABP3* was positively correlated with the right atrial diameter and the intra-atrial EMD, which are parameters of structural and electrical remodeling of the atrium. In contrast, there was no increase in the atrial content of DAG despite the increased atrial expression of *FABP3* and higher serum levels of FFA in our patients with chronic AF. Intriguingly, the atrial expression of a gene related to autophagosome maturation, *LC3*, was increased in the chronic AF patients, and autophagy-related genes including *LC3* were positively correlated with the atrial expression of *FABP3.* To the best of our knowledge, this is the first study showing that the atrial expression of *FABP3* gene is increased in association with autophagy-related genes in patients with chronic AF.

### The dysregulated fatty acid metabolism in the atrium of chronic AF patients

Compared to the SR patients, the chronic AF patients had higher serum FFA levels. The Cardiovascular Health Study has reported that an increase in the plasma levels of FFA by 200 μmol/L presents an 11% higher risk of AF occurrence even after adjustment for confounding risk factors including age, sex, race, physical activity, body mass index, coronary heart disease, congestive heart failure, smoking, alcohol use, log-C-reactive protein, diabetes mellitus, and hypertension in older adults [6]. Accordingly, elevated FFA levels can be an independent risk factor of AF.

The fatty acid-binding proteins (FABPs) reversibly bind to fatty acid and other lipophilic molecules. FABPs, which are located on the plasma membrane, facilitate fatty acid uptake into the cells, and intracellular FABPs transport fatty acid to other locations such as the nucleus and mitochondrion. Among the 10 isoforms of FABPs distributed in various tissues in mammals, FABP3 is most predominantly expressed in the heart [20]. Here, we observed that the gene expression of *FABP3* in the right atrial muscle was enhanced in chronic AF patients, which may indicate increased fatty acid uptake into the cells and increased intracellular fatty acid transport in the atrial muscle in chronic AF. In contrast, the gene expression of *CPT-1B* (which facilitates fatty acid transport across the outer mitochondrial membrane) and β-HAD activity (an enzymatic activity of mitochondrial fatty acid β-oxidation) in the atrium were comparable between our AF and SR groups.

### The association of the atrial expression of FABP3 with structural and electrical atrial remodeling

The results of our analyses revealed that the expression of *FABP3* in the right atrial muscle was positively correlated with the right atrium diameter and the intra-atrial EMD in the SR patients. The intra-atrial EMD is the time delay from the electrical activation to the actual motion of the atrial myocardium, and a delayed intra-atrial EMD indicates excitation-contraction uncoupling in the atrium. Prolonged intra-atrial EMD after cardioversion was reported to predict AF recurrence in patients with persistent AF, and histopathological changes characterized by myocardial fibrosis in the atrium appear to be a major determinant of the prolonged intra-atrial EMD [21].

Boldt et al. revealed that the atrial expression of collagen type I is enhanced in patients with lone AF, indicating that the occurrence of AF can directly increase the expression of collagen type I and cause myocardial fibrosis in the atrial muscle [22]. Although we did not conduct histopathological evaluations, previous reports and our present findings raise the possibility that the dysregulation of atrial fatty acid metabolism is linked to structural and electrical atrial remodeling, which may contribute to a future onset or recurrence of AF.

### Altered autophagy in the atrium in chronic AF

Garcia et al. first reported impaired cardiac autophagy characterized by reduced LC3 processing (i.e., a reduced protein expression of LC3BⅡ) with an accumulation of lipofuscin deposit — a potential trigger of AF — in the atrial myocardium in patients with post-operative AF [15]. A pair of studies have shown that in chronic AF patients, the cardiac autophagy characterized by an increased protein expression of LC3BⅡ is induced in the atrial myocardium in association with AMPK or endoplasmic reticulum (ER) stress [13, 14]. Our present findings demonstrated that the atrial gene expression of *LC3* was upregulated in the patients with chronic AF. Taking these results together, we speculate that altered cardiac autophagy in the atrium may be involved in the progression of AF.

### Implications of the association between the atrial expression of FABP3 and autophagy

The intracellular lipid content is generally deteremined by an imbalance between the uptake and the utilization of fatty acid, and thus the increased atrial expression of *FABP3* that we observed might contribute to the accumulation of lipids including DAG in the atrial myocardium in chronic AF. However, we did not detect an accumulation of DAG in the atrium in our patients with chronic AF as was reported in another study [23]. Autophagy has been shown to play a role in the regulation of fatty acid metabolism via the degradation of excessive intracellular lipids, termed “lipophagy” [12]. Our findings of an association between the atrial expression of *FABP3* gene and autophagy-related genes in chronic AF patients may support our hypothesis that in chronic AF, autophagy at least in part contributes to the prevention of the accumulation of toxic fatty acid metabolites via a degradation of intracellular lipids. Further research is necessary to clarify the mechanistic roles of cardiac autophagy in the atrium in AF progression.

### The difference in the atrial expression of FABP3 between post-operative AF and chronic AF

We observed that the atrial gene expression of *FABP3* was reduced in patients with post-operative AF in our prior study [16]; this is inconsistent with our present results regarding chronic AF patients. One of the possible explanations is a difference in pathophysiology between post-operative AF and chronic AF. It was demonstrated that in patients with metabolic syndrome, impairment in the mitochondrial respiratory capacity in the atrial tissues predicts the occurrence of post-operative AF [24]. In contrast, the mitochondrial respiratory capacity in the atrium was reported to be increased in chronic AF patients [25]. Taken together, these findings indicate that impaired energy metabolism (including reduced fatty acid utilization) in the atrial muscle might be a primary pathogenesis of post-operative AF, but in chronic AF, excessive fatty acid uptake into the atrial cells seems to play a crucial role in AF progression.

### Study limitations

Several limitations of this study should be addressed. First, most of the patients had valvular heart diseases, and the results of this study thus cannot be directly applied to patients with lone AF. Second, we were unable to perform western blotting to assess the protein levels of the autophagic marker due to the limited number of specimens. Therefore, based on our results alone we cannot definitively show whether autophagic flux is activated in the atrium. Finally, we cannot conclude that there is a causal relationship between fatty acid metabolism and autophagy in the atrium.

## Conclusions

Our study is the first to demonstrate that compared to patients with SR, the atrial expression of *FABP3* gene was upregulated in association with autophagy-related genes in patients with chronic AF. We also observed that the atrial gene expression of *FABP3* was related to structural and electrical remodeling in SR patients. Despite the increased atrial expression of *FABP3* with higher serum levels of FFA, atrial contents of DAG were not increased in patients with chronic AF. These findings provide new insights into the pathophysiology of chronic AF, and they suggest that dysregulated cardiac fatty acid metabolism might contribute to the progression of AF and induction of autophagy might have a cardioprotective effect against cardiac lipotoxicity in chronic AF.

## Acknowledgements

We thank Haruki Niwano for the technical assistance.

## References

1. Benjamin EJ, Wolf PA, D’Agostino RB, Silbershatz H, Kannel WB, Levy D. Impact of atrial fibrillation on the risk of death: the Framingham Heart Study. Circulation. 1998;98: 946–952.

2. Schnabel RB, Rienstra M, Sullivan LM, Sun JX, Moser CB, Levy D, et al. Risk assessment for incident heart failure in individuals with atrial fibrillation. Eur J Heart Fail. 2013;15: 843–849.

3. Wolf PA, Abbott RD, Kannel WB. Atrial fibrillation as an independent risk factor for stroke: the Framingham Study. Stroke. 1991;22: 983–988.

4. Wolf PA, Mitchell JB, Baker CS, Kannel WB, D’Agostino RB. Impact of atrial fibrillation on mortality, stroke, and medical costs. Arch Intern Med. 1998;158: 229–234.

5. Karam BS, Chavez-Moreno A, Koh W, Akar JG, Akar FG. Oxidative stress and inflammation as central mediators of atrial fibrillation in obesity and diabetes. Cardiovasc Diabetol. 2017;16: 120.

6. Khawaja O, Bartz TM, Ix JH, Heckbert SR, Kizer JR, Zieman SJ, et al. Plasma free fatty acids and risk of atrial fibrillation (from the Cardiovascular Health Study). Am J Cardiol. 2012;110: 212–216.

7. Lau DH, Nattel S, Kalman JM, Sanders P. Modifiable risk factors and atrial fibrillation. Circulation. 2017;136: 583–596.

8. Goette A, Honeycutt C, Langberg JJ. Electrical remodeling in atrial fibrillation. Time course and mechanisms. Circulation. 1996;94: 2968–2974.

9. Casaclang-Verzosa G, Gersh BJ, Tsang TS. Structural and functional remodeling of the left atrium: clinical and therapeutic implications for atrial fibrillation. J Am Coll Cardiol. 2008;51: 1–11.

10. Choi JY, Jung JM, Kwon DY, Park MH, Kim JH, Oh K, et al. Free fatty acid as an outcome predictor of atrial fibrillation-associated stroke. Ann Neurol. 2016;79: 317–325.

11. D’Souza K, Nzirorera C, Kienesberger PC. Lipid metabolism and signaling in cardiac lipotoxicity. Biochim Biophys Acta. 2016;1861: 1513–1524.

12. Singh R, Kaushik S, Wang Y, Xiang Y, Novak I, Komatsu M, et al. Autophagy regulates lipid metabolism. Nature. 2009;458: 1131–1135.

13. Yuan Y, Zhao J, Yan S, Wang D, Zhang S, Yun F, et al. Autophagy: a potential novel mechanistic contributor to atrial fibrillation. Int J Cardiol. 2014;172: 492–494.

14. Wiersma M, Meijering RAM, Qi XY, Zhang D, Liu T, Hoogstra-Berends F, et al. Endoplasmic reticulum stress is associated with autophagy and cardiomyocyte remodeling in experimental and human atrial fibrillation. J Am Heart Assoc. 2017;6: e006458.

15. Garcia L, Verdejo HE, Kuzmicic J, Zalaquett R, Gonzalez S, Lavandero S, et al. Impaired cardiac autophagy in patients developing postoperative atrial fibrillation. J Thorac Cardiovasc Surg. 2012;143: 451–459.

16. Shingu Y, Yokota T, Takada S, Niwano H, Ooka T, Katoh H, et al. Decreased gene expression of fatty acid binding protein 3 in the atrium of patients with new onset of atrial fibrillation in cardiac perioperative phase. J Cardiol. 2018;71: 65–70.

17. Xu ZX, Zhong JQ, Zhang W, Yue X, Rong B, Zhu Q, et al. Atrial conduction delay predicts atrial fibrillation in paroxysmal supraventricular tachycardia patients after radiofrequency catheter ablation. Ultrasound Med Biol. 2014;40: 1133–1137.

18. Zoghbi WA, Enriquez-Sarano M, Foster E, Grayburn PA, Kraft CD, Levine RA, et al. Recommendations for evaluation of the severity of native valvular regurgitation with two-dimensional and Doppler echocardiography. J Am Soc Echocardiogr. 2003;16: 777–802.

19. Takada S, Masaki Y, Kinugawa S, Matsumoto J, Furihata T, Mizushima W, et al. Dipeptidyl peptidase-4 inhibitor improved exercise capacity and mitochondrial biogenesis in mice with heart failure via activation of glucagon-like peptide-1 receptor signalling. Cardiovasc Res. 2016;111: 338–347.

20. Yamamoto T, Yamamoto A, Watanabe M, Matsuo T, Yamazaki N, Kataoka M, et al. Classification of FABP isoforms and tissues based on quantitative evaluation of transcript levels of these isoforms in various rat tissues. Biotechnol Lett. 2009;31: 1695–1701.

21. Ari H, Ari S, Akkaya M, Aydin C, Emlek N, Sarigul OY, et al. Predictive value of atrial electromechanical delay for atrial fibrillation recurrence. Cardiol J. 2013;20: 639–647.

22. Boldt A, Wetzel U, Lauschke J, Weigl J, Gummert J, Hindricks G, et al. Fibrosis in left atrial tissue of patients with atrial fibrillation with and without underlying mitral valve disease. Heart. 2004;90: 400–405.

23. Gizurarson S, Stahlman M, Jeppsson A, Shao Y, Redfors B, Bergfeldt L, et al. Atrial fibrillation in patients admitted to coronary care units in western Sweden - focus on obesity and lipotoxicity. J Electrocardiol. 2015;48: 853–860.

24. Montaigne D, Marechal X, Lefebvre P, Modine T, Fayad G, Dehondt H, et al. Mitochondrial dysfunction as an arrhythmogenic substrate: a translational proof-of-concept study in patients with metabolic syndrome in whom post-operative atrial fibrillation develops. J Am Coll Cardiol. 2013;62: 1466–1473.

25. Slagsvold KH, Johnsen AB, Rognmo O, Hoydal MA, Wisloff U, Wahba A. Mitochondrial respiration and microRNA expression in right and left atrium of patients with atrial fibrillation. Physiol Genomics. 2014;46: 505–511.

